# Peatland restoration can provide climate change mitigation over all time-scales: A UK case-study

**DOI:** 10.1101/2024.12.10.627721

**Authors:** Xiao Zhang, Minna Ots, Eleanor Johnston, Kathryn Brown, Nigel Doar, John Lynch

## Abstract

Peatlands provide one of the largest terrestrial carbon stocks in the UK. However, a large proportion of peatlands are drained for peat extraction, agriculture and other uses, turning them into a major source of the UK’s land use greenhouse gas (GHG) emissions. Successful restoration can ultimately return peatlands into carbon sinks. However, rewetting - the primary step in peatland restoration - can reduce CO_2_ emissions while increasing CH_4_ emissions. This may result in little overall climate benefit, or even increased warming for several years post peatland restoration, as CH_4_ is a short-lived but strong GHG, and may overpower the reduction in CO_2_. Such consequences are rarely explored in detail, since most studies are based on comparing total CO_2_-equivalent emissions pre- and post-restoration using the 100-year Global Warming Potential (GWP100), which can fail to reveal the full dynamics. We evaluated the emissions and resultant climate impacts from peatland restoration using data from The Wildlife Trusts, a federation of UK-based conservation charities, as a case-study. The total emissions of each restoration stage were estimated by multiplying peatland areas under restoration with up-to-date UK emission factors (EF), then compared under multiple pulse emission metrics (GWP100, GWP20, GTP100) to indicate the impacts over a range of time-horizons, and GWP* to reveal the varying warming impacts over time. We also used Monte-Carlo Simulation to investigate the uncertainties in total emissions drawing from EF ranges. We found that the restoration so far has provided large emission reductions under all metrics, even considering the uncertainties. Increased CH_4_ is unlikely to cause extra warming in the extremely near-term (<20 years), and if the peatlands are maintained in their rewetted states, they can contribute to net-cooling in the long term. There is less certainty over the climate benefits of further restoration, from rewetted to “near-natural” states, especially in the shorter term, but we argue that any risks are low, while this continued restoration will provide further ecological benefits and support biodiversity. Our study lends further support for peatland restoration in the UK and other regions with similar habitats, and provides insight into the climate roles of peatlands more broadly.

## Introduction

An important component of climate change mitigation can be provided by nature-based approaches: conservation, restoration and improved management of ecosystems so that greenhouse gas (GHG) emissions are reduced and/or carbon is sequestered in habitats (Seddon et al., 2020; Girardin et al., 2021). Peatland conservation and restoration is a particularly important nature-based climate solution. Peatlands are the largest terrestrial carbon stock but widespread peatland degradation means that significant amounts of this carbon are being lost to the atmosphere, making them a major emission source (Strack et al., 2022). Peatland restoration can reduce these carbon losses, and instead result in peatland systems providing ongoing removal of atmospheric carbon dioxide (CO_2_), in addition to other ecosystem services, if in good ecological conditions (Page & Baird, 2016). However, the full climate impact of peatlands, and the extent to which restoration provides climate change mitigation, is complicated by the substantial methane (CH_4_) emissions associated with wetland habitats.

Natural peatlands (or managed peatlands that are functionally similar enough to result in the same dynamics) capture CO_2_ by slowing down aerobic respiration and preventing dead plant materials being fully decomposed, and emit CH_4_ through anaerobic respiration (Drösler et al., 2008). These occur as peatland soils are sufficiently waterlogged to create anoxic conditions. The overall effect of waterlogging is a reduction in decomposition of organic matter, and hence peatlands act as a net carbon sink with photosynthetically fixed carbon accumulating over time (Page and Baird, 2016). When peatlands are drained and exploited, the water table decreases, leaving the carbon-rich soil exposed. The carbon is released via a large amount of direct CO_2_ emission from oxidation of the organic materials, as well as running-off through dissolved organic carbon (DOC) and particulate organic carbon (POC) in the water (IPCC, 2014). The aerobic condition also favours nitrification and thus increases nitrous oxide (N_2_O) emissions (Martikainen et al., 1993). While the direct CH_4_ emission is suppressed, a small amount of CH_4_ is produced by waterlogged drainage ditches (Peacock et al., 2021). Overall, degraded peatlands have higher CO_2_ and N_2_O emissions and lower CH_4_ emissions compared to the natural ones. Rewetting, as a reverse process of drainage, has the dual effect of CO_2_ uptake and CH_4_ emissions, and a review based on the northern peatlands showed rewetted peatlands on average have 46% more CH_4_ emissions than the pristine ones (Abdalla et al., 2016). Since CH_4_ is a more powerful GHG than CO_2_, concerns have been raised that rewetted peatlands may have a net warming impact, even if the site accumulates carbon (Neubauer and Megonigal, 2015; Ma, Creed and Badiou, 2024).

Understanding the net climate impact of peatlands is not straightforward, as the different GHGs associated with peatlands have markedly different dynamics. CH_4_ is a powerful but short-lived GHG whose instantaneous warming effect is more than 100 times stronger than CO_2_ (Dessus, Laponche & Le Treut, 2008) but its average atmospheric lifetime is only around a decade, with CH_4_ oxidized by hydroxy radicals in the atmosphere. CO_2_ is not chemically active in the atmosphere, and its anthropogenic emissions are generally considered to persist in the atmosphere for thousands of years. Continued emissions of the two gases therefore result in different dynamics: “short-lived” CH_4_ has a strong warming effect in the decades immediately after emission, but which then reduces over time as the emission decays, while CO_2_ emission has longer-term impacts which accumulate over time as long as emissions continue (Pierrehumbert, 2014). N_2_O also has a much stronger climate impact than CO_2_ per unit mass, and has an average atmospheric lifetime of just over 100 year. Thus, N_2_O acts cumulatively, akin to CO_2_, over periods of several decades, but on multi-centennial timescales its impacts also depend more on the continued flow of emissions, rather than the cumulative total.

Typically, non-CO_2_ GHGs such as CH_4_ and N_2_O are reported and assessed in the form of CO_2_-equivalent (CO_2_e) emissions, converted using a weighting factor. The most commonly used CO_2_ equivalence metric is the 100-year Global Warming Potential (GWP100), which is based on the relative radiative impact of pulse emissions of a gas averaged in 100 years. It has been argued that this may overlook the dynamics of shorter-vs longer-lived climate pollutants as described above (McAuliffe et al., 2023; Cherubini et al., 2016; Lynch et al., 2020), and the metric is ill-suited to understanding the climatic role of peatland emissions (Neubauer and Megonigal, 2015). The Intergovernmental Panel on Climate Change (IPCC, 2023) suggest that the choice of emission metrics should depend on the purpose of evaluation, and report multiple possible emission metrics that could provide insight into different dynamics and/or over different timeframes.

A key practical question that emerges is whether peatland restoration is a viable and worthwhile component of climate change mitigation. Rewetting is the most effective method to stop the oxidation process, and the first step to restoring the species composition (Beyer et al., 2021), However, the temporal dynamics in CH_4_-induced warming effect have raised the concern that, especially at the beginning of the restoration process, the temperature increase resulting from elevated CH_4_ may mean restoration *increases* overall warming, even if the peatland shifts from CO_2_ source to sink (Bridgham et al., 2006; Li et al., 2022). To address this concern, here we provide a case-study exploring pre- and post-restoration GHG emissions as reported under a range of emission metrics, with a particular focus on how different time-horizons influence evaluation, including GWP*, a novel approach to calculating ‘equivalent emissions’ that can illustrate changes over time (Smith, Cain & Allen, 2021; Cain et al., 2019). Assessing how changes in peatland GHG fluxes resulting from restoration impact warming over time will help increase the confidence in planning whether and where restoration should be encouraged.

We use peatland restoration scenarios in the United Kingdom (UK), building on real-life restoration examples and plans from The Wildlife Trusts in the UK. The Wildlife Trusts is a federation of conservation charities composed of 46 local Wildlife Trusts and a central charity, the Royal Society of Wildlife Trusts. Together, the Trusts undertake environmental management and conservation in local sites across the UK and also advocate for nature and climate action at a national and global level. The UK also provides an interesting and important case-study location building on prior research on northern peatlands (e.g. Harris et al., 2022; Qiu et al., 2022). 12% of the UK land area is covered by peatland (Trenbirth & Dutton, 2019), which has high carbon storage and supports unique ecosystems (JNCC, 2011). However, nearly 80% of UK peatlands are degraded due to draining for agriculture, forestry or peat extraction, and have become one of the largest land use related emission sources (Trenbirth & Dutton, 2019; Young et al., 2023), estimated to emit around 23 Mt CO_2_e yearly (Evans et al., 2017). Consequently, peatland restoration also features prominently in the UK’s land policy expectations to help deliver “net-zero” emissions by 2050 (Committee on Climate Change, 2020), and recent work has updated UK peatland emission factors (EF) to reflect the contemporary scientific understanding, and facilitate the incorporation of peatland restoration in the National Inventory Report of annual emissions and removals (Evans et al, 2023). However, these standard approaches and previous UK studies all, to our knowledge, assess net climate impacts using GWP100. This study addresses the gap in the assessment of warming under various time scales, better revealing the climate benefits of recent and planned peatland restoration in the UK, and contributing to discussions over how to conceptualise and report the net impact of peatland emissions more broadly.

## Methods

### Peatland restoration by The Wildlife Trusts

We use peatland restoration data from The Wildlife Trusts as a case study representing realistic restoration pathways, at a scale relevant for a range of national/sub-national decision making and emission inventory purposes. The Wildlife Trusts have been leading peatland restoration projects across the UK and gathering information on their restoration progress. The data has a wide coverage of England, Northern Ireland, Wales and Isle of Man and were collected from the individual Trusts in early summer 2024. The data records areas of the restored peatland sites, including habitat conditions during the restoration process that are reported in 3 generalised stages: initial conditions, current status and restoration target. The sites with no area or habitat conditions in any of the restoration stages were omitted, leaving a total of 57,367.5 hectares of peatlands under restoration being assessed in this case study.

In the data supplied from individual Trusts, the habitat types and conditions were recorded using similar typologies as reported in the source of the EFs (Evans et al., 2023; Table 1), hence correspond with the contemporary EFs. If habitat data were not already provided in one of these categories, the closest alignment from the description given was used. In addition, if the habitat types are unclear (e.g. reported simply as “modified drained”), we treated them as the possible habitat type with the highest EFs (e.g. modified bog – eroding (drained)). The two habitat categories not in Table 1, rewetted modified fen and forested peatland, were assumed to have the same EFs as near-natural fens and modified bog – eroding (drained), respectively. For the sites that contain multiple peatland habitat types in the same stage, we assumed the peatland is a mosaic of those habitat types, and each habitat type makes up an equal proportion of the area, if they did not have assigned proportions (i.e., if a site is described as “cropland - wasted (peat < 40 cm); drained - conifer plantation”, we assumed half the site is cropland, and half is conifer plantation). For the convenience of scenario design and the uncertainty analysis (see the section *Uncertainty analysis: Monte-Carlo Simulation*), we split these mosaics into individual sub-sites that contain only one habitat type, whose areas were evenly distributed over the total area of the mosaic habitat.

**Table 1.** Peatland condition categories, following Evans et al. (2023).

| Category |
| --- |
| Near-natural bog |
| Near-natural fen |
| Rewetted bog |
| Rewetted fen |
| Rewetted modified bog |
| Modified bog – grass/heather (drained) |
| Modified bog – grass/heather (undrained) |
| Modified bog – eroding (drained) |
| Modified bog – eroding (undrained) |
| Extracted – domestic (drained) |
| Extracted – industrial (drained) |
| Grassland – extensive (drained) |
| Grassland – intensive (drained) |
| Cropland (peat > 40 cm) (drained) |
| Cropland - wasted (peat < 40 cm) (drained) |

The initial conditions were primarily drained or otherwise degraded peatlands, with the first stage in restoration mainly focused on rewetting. The subsequent restoration stage, from current to target conditions, is more focused on restoring rewetted habitats to near-natural conditions. To illustrate, a typical evolution of a given site could be progressing across condition categories in Table 1 from extracted (industrial), to rewetted modified bog, to near-natural bog.

In general, the current conditions reported the habitat status in 2024, with individual sites showing a range of different starting years and planned ending years of restoration. To simplify our scenario design and interpretation of GHG impacts over time (discussed further below), we assume that the restoration transitions all occur in discrete stages at the same time across the entire area, and we report emission sums across the entire habitat area at each stage.

### Estimating greenhouse gas emissions

We used the EFs from Evans et al. (2023) to calculate the GHG emissions from The Wildlife Trusts’ restored peatlands. For many research and reporting purposes, particularly for official GHG emission inventories (National Inventory Reports as submitted to the United Nations Framework Convention on Climate Change; UNFCCC), it is standard practice to estimate GHG emissions by recording “activity data”, such as quantity of fossil fuels combusted or area of a given habitat, that is multiplied by EFs to estimate the corresponding GHG fluxes associated with that activity.

Evans et al. (2023) updated the Tier 2 (regionally specific, c.f. Tier 1 means global) EFs for UK peatland habitats, using the field emission data from only the UK or regions with similar climate and soil conditions. It provides the most recent and comprehensive classification of the UK’s peatland habitats and their EFs breaking down to individual gas emission pathways, which includes the direct emission of CO_2_, CH_4_ and N_2_O, as well as DOC, POC and ditch CH_4_. The EFs of direct CO_2_, CH_4_ and N_2_O also have associated standard errors (SE) and 5% and 95% confidence intervals (CI). We obtained the emissions estimations of each restoration stage by summing the product of mean per-gas EFs and areas of all the individual sites. Tracking changes individually for CO_2_, CH_4_ and N_2_O allow us to keep full details and enable exploration of gas-specific impacts over time, before also reporting as CO_2_-equivalent aggregations (see sections below on *CO_2_-equivalent emissions* and *‘warming equivalent’ emissions)*.

### CO_2_-equivalent (CO_2_e) emissions

As outlined in the introduction, non-CO_2_ GHGs are typically reported as ‘CO_2_-equivalent (CO_2_e) emissions’, and activities that emit multiple gases, all gases are aggregated as a total CO_2_e emission. GWP100 is the most commonly used emission metric, including for government and international reporting obligations (Dessus, Laponche & Le Treut, 2008). It shows the radiative forcing (change in atmospheric energy budget) resulting from a pulse emission of a given gas, relative to CO_2_ and averaged over a time period of 100 years. GWP100 reports CH_4_ as a moderately strong GHG, while the relative valuation of CH_4_ can be vastly different depending on the timescale and metric (Table 2). Therefore, we also used GWP20, which is the Global Warming Potential averaged over 20 years, to highlight the short-term effects of emissions, as it has a much greater valuation for CH_4_; and GTP100, the Global Temperature change Potential after 100 years which reflects relative contribution to temperature increases *after* 100 years, to show the longer-term impacts effect. The use of GWP20 and GTP100 to illustrate shorter- and longer-time impacts alongside GWP100 has been recommended under Life Cycle Assessment frameworks (Cherubini et al., 2016). Note that all of these metrics are based on the behaviours of individual pulse emissions. The CO_2_e emissions were calculated for all three of these emission metrics by multiplying the respective individual gas quantities by the respective values taken from the IPCC 6^th^ Assessment Report (AR6) as shown in in Table 2.

**Table 2.** Total emissions per year from The Wildlife Trusts’ restored peatlands for each GHG emission pathway, calculated using the central value of EFs.

|  | **GWP100** | **GWP20** | **GTP100** |
| --- | --- | --- | --- |
| **Non-fossil CH<sub>4</sub>** | 27.0 | 79.7 | 4.7 |
| **N<sub>2</sub>O** | 273 | 273 | 233 |

**Table 2.**
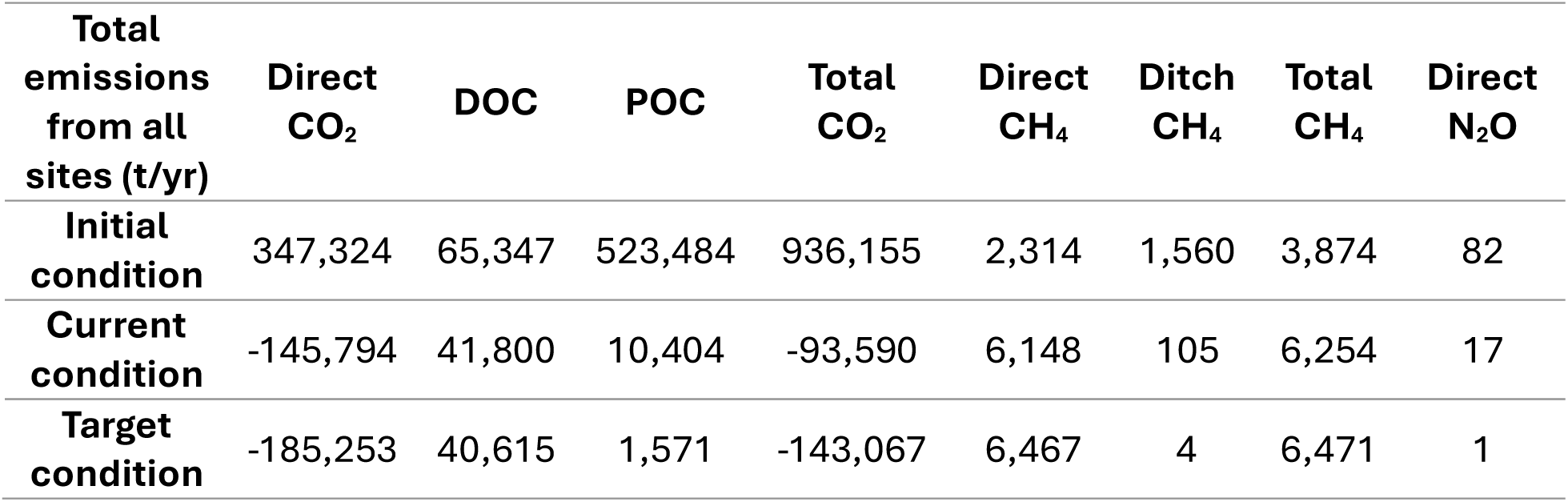
IPCC 6^th^ Assessment Report metric values of CO_2_ equivalent.

| **Total emissions from all sites (t/yr)** | **Direct CO<sub>2</sub>** | **DOC** | **POC** | **Total CO<sub>2</sub>** | **Direct CH<sub>4</sub>** | **Ditch CH<sub>4</sub>** | **Total CH<sub>4</sub>** | **Direct N<sub>2</sub>O** |
| --- | --- | --- | --- | --- | --- | --- | --- | --- |
| **Initial condition** | 347,324 | 65,347 | 523,484 | 936,155 | 2,314 | 1,560 | 3,874 | 82 |
| **Current condition** | -145,794 | 41,800 | 10,404 | -93,590 | 6,148 | 105 | 6,254 | 17 |
| **Target condition** | -185,253 | 40,615 | 1,571 | -143,067 | 6,467 | 4 | 6,471 | 1 |

In addition to the GWP100, GWP20 and GTP100 aggregated total emissions across all sites at each restoration stage (“initial”, “current”, “target”), we also calculated the difference in emissions between the “initial” and “current” conditions (emissions savings so far), and between the “current” and “target” conditions (further emissions savings).

### Alternative approach to report emissions as ‘warming equivalents’ (we)

While the multiple metrics described above can give some insight into temporal dynamics, they do not reveal the full evolution of warming contribution over time. As noted in the previous section, the metrics reflect the impact of pulse emissions, whilst most peatland emissions will be sustained to some degree. They also have limited capacity to evaluate when the contribution of peatlands may shift from warming to cooling effects (Neubauer, 2014).

To investigate the evolution of the impact of different gases, we used a relatively novel “CO_2_ warming equivalent (CO_2_-we)” emission approach, GWP* (Allen et al., 2016; Cain et al., 2019; Smith, Cain & Allen, 2021) (Equation 1). It effectively reports individual CH_4_ emissions as a very large CO_2_ release, followed by an automatic CO_2_ removal after 20 years, reflecting the strong but temporary impact of CH_4_ emissions. GWP* describes emission scenarios as “warming-equivalents”, providing a direct approximation of the warming that results over time from these emissions, in line with the warming dynamics that would be observed if the “CO_2_-we” was emitted as CO_2_ itself (Lynch et al., 2020). Furthermore, cumulative CO_2_-we emissions have a direct linear correspondence with the associated temperature increase, and so can be interpreted as directly reveal temperature dynamics over time: this is because CO_2_ itself shows a linear relationship between total warming and cumulative emissions, but the ‘conventional’ CO_2_e under any pulse emission metrics does not work in this manner (Allen et al., 2022).

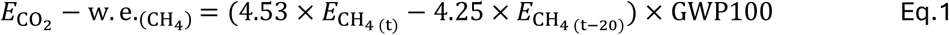

We used GWP* to show the climate impact over time and model different restoration scenarios. We created 3 arbitrary scenarios with the habitats in their “initial conditions” and associated emissions beginning from year 0, and illustrated their cumulative CO_2_-we over 200 years, where CH_4_ was converted using GWP* (Eq.1), N_2_O was converted using GWP100 and CO_2_ added without conversion (Note that with an average lifetime of shortly over 100-years, N_2_O can also be treated as having a cumulative impact reflected by its GWP100 CO_2_-equivalence for periods up to a couple of centuries, as discussed in Lynch et al, 2020). The emission rates are obtained from the *Estimating greenhouse gas emissions* section.

In Business As Usual (BAU) scenario, we assumed all habitats remained in their “initial conditions”. In our Average 2-step Restoration (A2R) scenario, all habitats were restored to “current conditions” on year 75, and were then restored to “target conditions” on year 100 and remained to year 200. To compare the climate impacts of restoring to rewetted conditions and to near-natural conditions, in the Average 1^st^-step-only Restoration (A1R) scenario, all habitats were only restored to “current conditions” on year 75 and remained till year 200. While arbitrary, these specific times were chosen so that a) there is an initial period revealing the impact of the initial peatland emissions, reflecting that much of the UK’s peatland is already degraded and has been for some time (as can be inferred in the results below, the length of this initial period changes the extent of warming accrued while degraded, but does not affect the general dynamics or relative outcomes of future restoration); b) there may be challenges in restoring biodiversity and wider ecosystem function that takes longer than initial re-wetting, with 25-year period between the “current” and “target condition” selected to help separate out the impacts of restoration vs effects of CH_4_ over time (noting the dependency of GWP* on 20-year emission periods); and c) to highlight the ongoing, longer-term climate impacts of sustaining emissions in either state.

In the BAU, A2R and A1R scenarios the total emission rates were all calculated using the mean EFs. To investigate the potential of the second step restoration having a much worse-than-average outcome within the range of EF uncertainties we created an Outlier 2-step Restoration (O2R) scenario, where the habitat condition changes still happen on year 75 and 100 as described, but the emission rate of the “current conditions” was calculated using the 5% CI EFs, and the emission rate of the “targeting conditions” was calculated using the upper 95% CI EFs, representing a more positive situation for the “current condition” and a more negative outcome for the “targeted condition”.

### Uncertainty analysis: Monte-Carlo Simulation

Using the uncertainties of direct CO_2_, CH_4_ and N_2_O EFs in Evans et al. (2023), we conducted the uncertainty analysis for the total emissions of each restoration stage as reported under the 3 pulse emission metrics. Since the shapes of the EF distributions were not described, if the distances of 5% and 95% CI to the means were the same (i.e. they were symmetrical), we assumed a normal distribution; if not (i.e. if the 5% and 95% CI were asymmetrical about the mean) then we assumed log normal distribution (Milne et al., 2014). The EFs of DOC, POC and ditch CH_4_ have no uncertainty information, so we used the means for all iterations. We also assumed there is no uncertainty in the land areas or condition scores received from the Trusts.

Monte-Carlo simulation was undertaken by randomly picking EFs from their distributions for calculating the total emissions in each iteration. Within each iteration, EFs were selected independently for each site, allowing sites of the same habitat category to have distinct EFs. This approach realistically reflects site-specific emissions variabilities. Our preliminary tests showed that 100,000 iterations produced reliable distributions of the total emissions, with further runs having no major impact on distribution shape or other parameters. Since the simulation outcomes for all stages have the shape of normal distribution (Figure 3), the central values of the total emissions are the means of all iterations. We also calculated the SE and 5% and 95% CI (Table S1). To explore the implications of uncertainties across restoration stages and determine the likelihood that restoration reduces or increases aggregated CO_2_-equivalent emissions, we compared the total emissions of different restoration stages within a pairwise set of Monte-Carlo iterations, and summarized the percentage of 100,000 iterations that have lower/higher emissions after restoration (method adapted following Flysjö et al., 2011).

Due to the large number of simulations, we did not perform statistical tests comparing central tendency (i.e. t-tests or ANOVA comparing means), as central tendency can be reliably estimated when there is a large number of samples, but it does not provide insight into the underlying environmental considerations.

The analyses were performed in Microsoft Excel and R version 4.3.0 (R Core Team, 2023).

## Results

### Individual GHG emissions

The estimated emissions, using the mean EFs, reduced substantially from most of the pathways after restoration (Table 2). At the initial stage, CO_2_ was the major source of peatland GHG emissions, with a moderate amount of CH_4_ emissions and small amount of N_2_O. Among all sources of CO_2_ emissions, POC caused 523 Kt/yr emissions and had the greatest contribution. After restoring to the “current stage”, emissions from POC and direct CO_2_ dropped hugely, and the sites became CO_2_ sinks. The CO_2_ sinks would further grow when being restored to “target conditions”. Similarly, the direct N_2_O emissions have also been decreasing, and the final restored habitats would have an emission rate of 1 t/yr, becoming nearly N_2_O neutral. CH_4_ emissions follow an opposite trend. The total CH_4_ emissions had a large leap from “initial” to “current conditions”, and a small further increase when turned into “target conditions”. Although there is a reduction in CH_4_ emissions from drainage ditches after restoration, direct CH_4_ emissions keep increasing significantly and make total CH_4_ emissions increase.

On average, restoring to “current conditions” has reduced direct CO_2_ and POC emissions by 8.60 t/ha/yr and 8.94 t/ha/yr respectively, while the increase in direct CH_4_ emissions is 0.067 t/ha/yr.

### CO_2_-equivalent emissions

The standard metric GWP100 shows that there has been approximately 980 KtCO_2_e/yr emissions reduction achieved by restoring the sites to “current condition”. If they are restored to their “target conditions”, there will be a small amount of further emissions savings (Table 3).

**Table 3.** The total CO_2_-equivalent emissions per year of each stage of The Wildlife Trusts restored peatland habitats and the emissions savings of each step, converted using the IPCC metric values in Table 1.

| **Total emissions**<br>**(KtCO<sub>2</sub>e/yr)** | **GWP100** | **GWP20** | **GTP100** |
| --- | --- | --- | --- |
| **Initial condition** | 1,063 | 1,267 | 973 |
| **Current condition** | 80 | 410 | -60 |
| **Target condition** | 32 | 373 | -112 |
| **Emissions saving**<br>**so far** | 983 | 858 | 1,034 |
| **Future emissions**<br>**saving** | 48 | 37 | 52 |

Using GWP20, the total emissions in CO_2_ equivalent are much higher than in GWP100 under all 3 stages as the converting factor for CH_4_ is much higher (Table 1). The emissions savings from restoration so far is lower than in GWP100 but still prominent (858 eq KtCO_2_/yr) (Table 3).

On the other hand, the long-term impact of restoration illustrated using GTP100 showed the greatest benefit, giving the highest emission savings both so far, and in the future if restored to target conditions, all three metrics (Table 3). Under GTP100, the reported CH_4_ emissions are valued sufficiently low that aggregated emissions are dominated by CO_2_ removals, with the first restoration step (up to present day) already resulting in net-negative emissions, which are nearly doubled in the second restoration step (future restoration).

### Warming over time: GWP*

In the Business As Usual (BAU) scenario, the cumulative CO_2_ and N_2_O emissions kept rising at a steady rate through the whole time period, while cumulative CH_4_ warming equivalent increased faster in the first 20 years. CO_2_ had the greatest contribution to the total cumulative CO_2_-we emissions, and makes up an increasingly large share of cumulative emissions (hence net warming) the longer that all emissions continue (Figure 1A).

**Figure 1.**
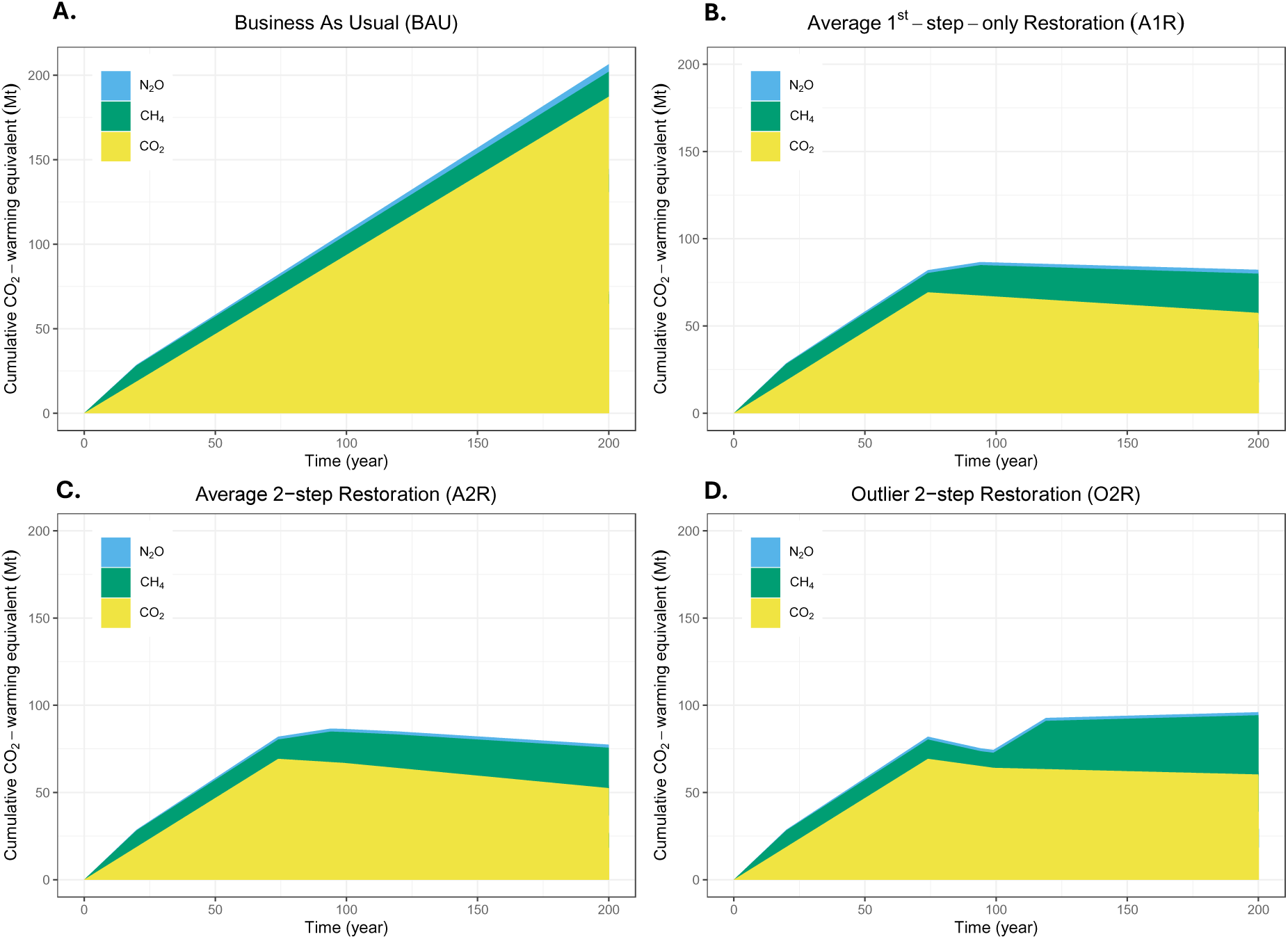
Cumulative ‘CO_2_-warming equivalent’ emissions with the separation of each gas, where the warming effect of CH_4_ is assessed using GWP*. Scenarios assume zero emissions initially as arbitrary starting point of degraded peatlands and generation of emissions. In the **Business As Usual (BAU)** scenario (A.), all sites remain in the “initial condition” for 200 years. In the **Average 1^st^-step-only Restoration (A1R)** scenario (B.), all sites only change from “initial conditions” to “current conditions” on year 75. In the **Average 2-step Restoration (A2R)** scenario (C.), all sites change from “initial conditions” to “current conditions” on year 75, and then change to “target conditions” on year 100. In the BAU, A1R and A2R scenarios, all of the emission rates were calculated using the central values of EFs. In the **Outlier 2-step Restoration (O2R)** scenario (D.), all sites also experience the same condition changes as in the A2R scenario, but the emission rate of the “current conditions” was calculated using the lower 95% CI EFs, and the emission rate of the “target conditions” was calculated using the upper 95% CI EFs.

In the Average 2-step Restoration (A2R) scenario, cumulative CO_2_ started to decrease right after the first restoration, as the peatland switches from a CO_2_ source to sink, while CH_4_ warming effect rose at a high rate between year 75 and 95. The cumulative impact of N_2_O warming effect is nearly negligible compared to the other two gases, even at the end of the 200-year period (Figure 1C). The total cumulative warming equivalent started to grow much slower than in BAU immediately after the first restoration, and started to mildly decrease on year 95 (Figure 2), which was an effect from natural CH_4_ removal after being increased by the first restoration for 20 years. Despite having greater CH_4_-induced warming than in the “initial conditions”, the CO_2_ removals are great enough to lead to a declining temperature. Comparing with the Average 1^st^-step-only Restoration (A1R) scenario, the second restoration on year 100 contributed to a small amount of further net cooling (Figure 2), as there was a higher rate of CO_2_ removal, and only a small increase of CH_4_ emission rate comparing with the previous stage.

**Figure 2.**
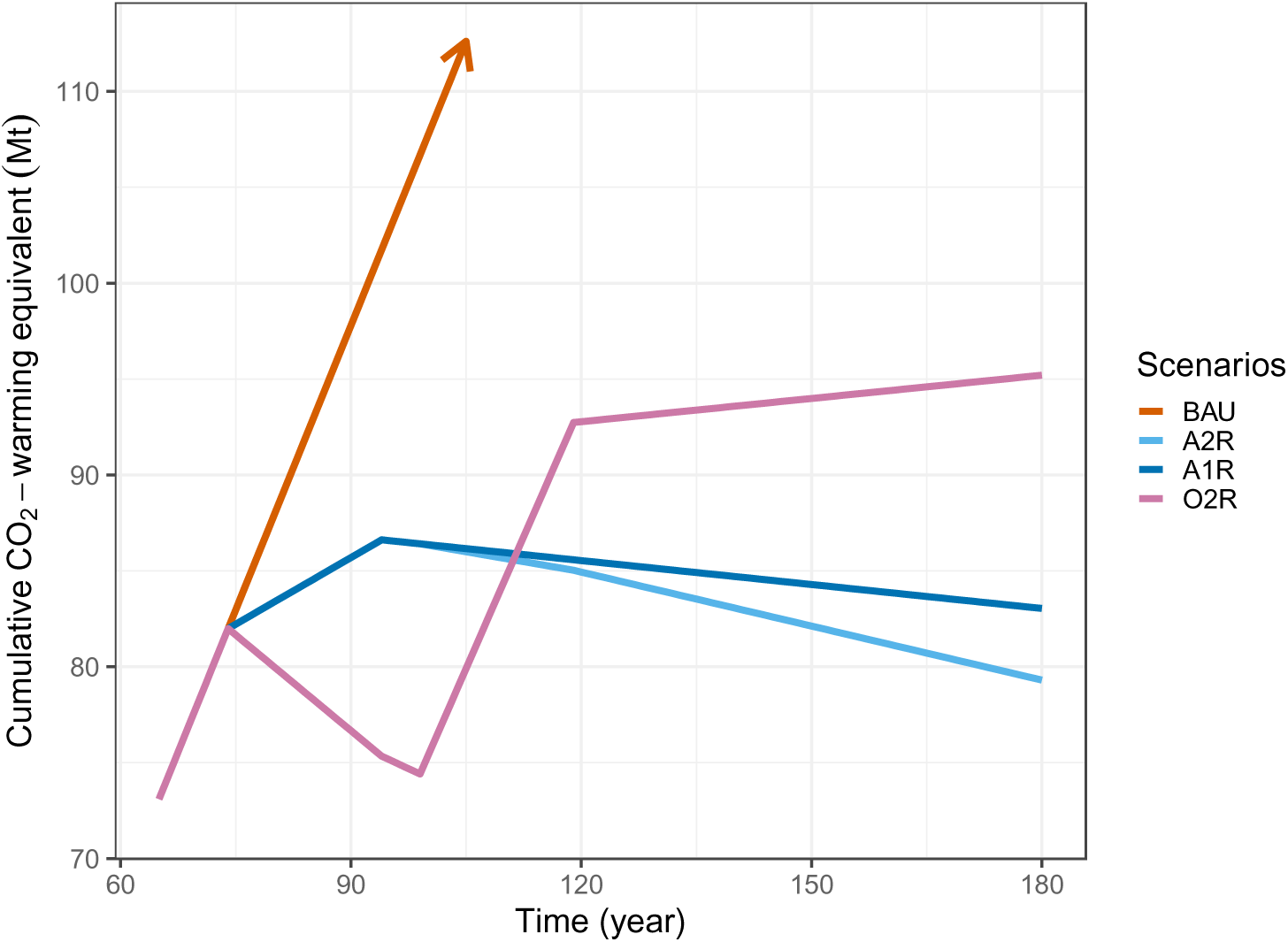
Cumulative total ‘CO_2_-warming equivalent’ emissions of the restoration scenarios designed in Figure 1. Scenarios are: Business As Usual (BAU); Average 1-step-only Restoration (A1R); Average 2-step Restoration (A2R); and Outlier 2-step Restoration (O2R). See Figure legend 1 or text for full explanation of scenarios.

In the Outlier 2-step Restoration (O2R) scenario, the CO_2_ removal rate for the optimal “current conditions” (year 75-100) was greater than the worst-case “target conditions” (year 100-200). The trend of the N_2_O warming effect remained the same as in the A2R scenario. CH_4_ emissions experienced the most dramatic changes, and the change in cumulative total warming equivalent was mostly determined by CH_4_ balance (Figure 1D). Compared to A2R, the CH_4_ warming effect remained low after the first restoration, but increased rapidly after the second restoration, eased at year 120 and increased only mildly afterwards. The changes in the CH_4_ warming effect caused the net cumulative warming effect to decline significantly in year 75-100, then rapidly rise in year 100-120, and grow gently afterwards. Despite the rapid changes after restoration, the total cumulative balance and its rate of increase are both much lower than in the BAU scenario.

### Monte-Carlo Simulation Uncertainty analysis

The mean values of the total emissions from the Monte-Carlo simulation are similar to the estimation using central value EFs (Table S1, Table 3). The estimations using GWP20 have wider distributions than those using GWP100 and GTP100, but due to the large sample size and peaked distributions (resulting from samples reflecting aggregate totals of multiple sampling selections), all estimations have very small SE and CI (Table S1).

The total emissions of the “initial condition” are much higher than restored conditions and their distributions have no overlap in all 3 metrics (Figure 3), showing that restoration so far has significantly reduced emissions by a large amount, regardless of uncertainties. When comparing “current” and “target conditions”, in GWP100 and GTP100, 70.8% and 84.5% of the simulations reduce emissions after the second restoration respectively, while the figure for GWP20 is only 57.8%: i.e., under GWP20, the second restoration step towards ‘near natural’ conditions has a 42.2% chance of *increasing* aggregate CO_2_e emissions.

**Figure 3.**
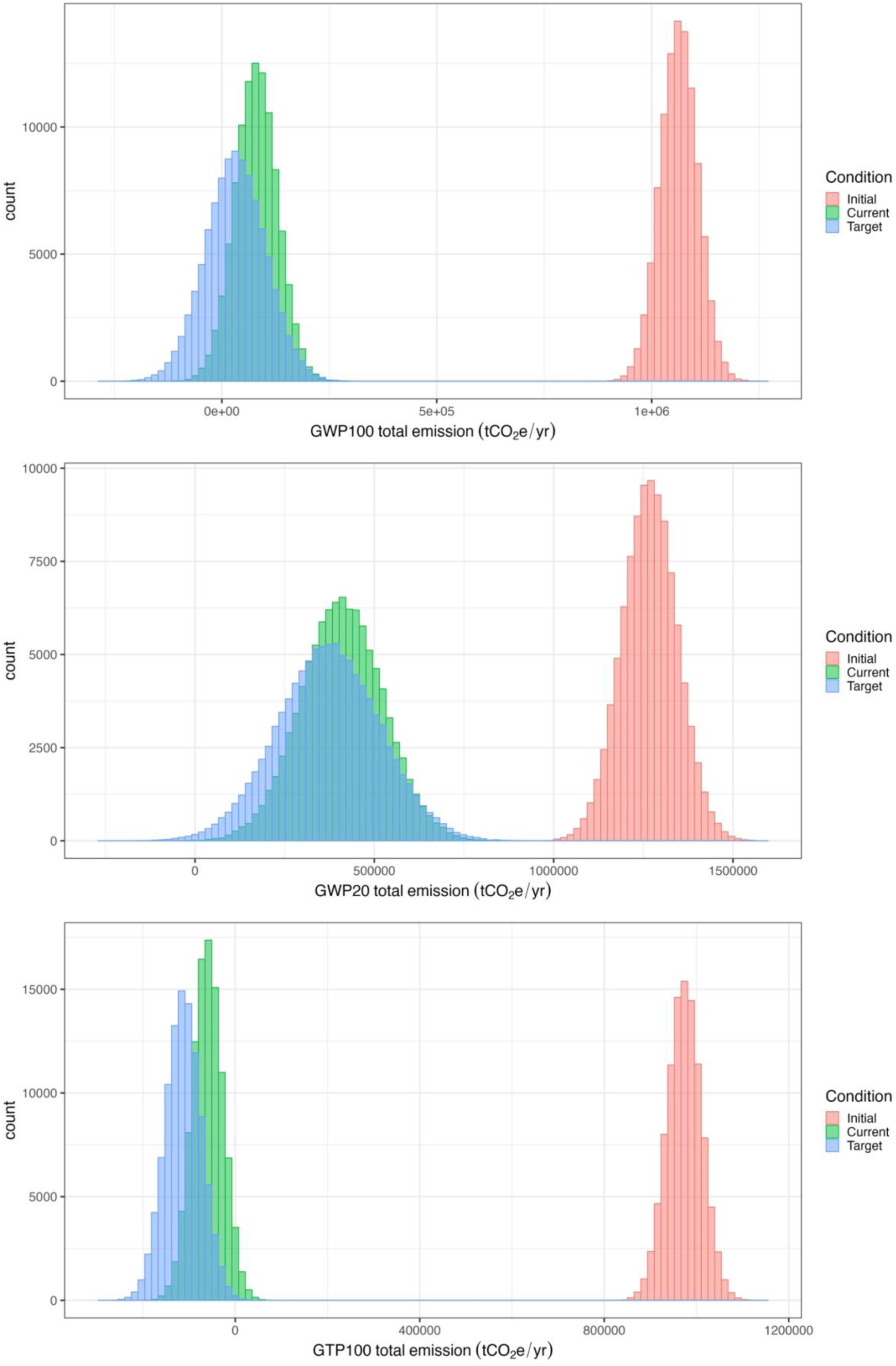
Histograms showing the distributions of total emissions each year for all the sites in their “initial”, “current” and “target” conditions respectively obtained by Monte-Carlo simulation, in GWP100 (top), GWP20 (middle) and GTP100 (bottom).

## Discussion

### Impact of peatland restoration on GHG fluxes

When assessing each GHG independently, we expect that the peatland restoration activities undertaken by The Wildlife Trusts have resulted in decreased CO_2_ losses – with restored peatlands switching from emitting to removing atmospheric CO_2_ – along with reduced N_2_O.

Further restoration towards near-natural conditions would be expected to result in ever greater CO_2_ removals and further N_2_O reductions, but a small additional increase in CH_4_. These expectations stem primarily from the EFs associated with different peatland conditions in the latest methodological update for the UK (Evans et al 2023). These are scientifically grounded and will underpin future reporting for official inventory purposes (and hence whether and how peatland restoration contributes to a ‘net-zero’ UK, from an emissions accounting perspective), making them important for exploring in modelling studies such as this. However, there is still an imperative for future work to continue assessing GHG fluxes via *in situ* monitoring to confirm and/or update the EFs in the future and continue to assess their suitability for a diversity of peatland habitats and restoration pathways, which should be kept in mind while interpreting the EF-based emissions estimations.

The UK specific EFs from Evans et al. (2023) and hence the anticipated gas-specific trends observed here reflect the impact of rewetting on emissions, with higher water tables reducing CO_2_ but increasing CH_4_ (Evans et al., 2021), and are comparable to other studies. A meta-analysis based on the primary measurement of direct CO_2_ and CH_4_ emissions changes after rewetting had the same overall conclusions, where the reduction in CO_2_ emissions and increase in CH_4_ emissions were both significant, while the magnitudes of changes were slightly different (Darusman et al., 2023). Our estimation of the amount of CO_2_ decrease and CH_4_ increase are both higher, as Darusman et al. (2023) found the CO_2_ emissions reduced by −1.43 ± 0.35 Mg CO_2_–C ha^−1^yr^−1^, and increased CH_4_ emissions by 0.033 ± 0.003 Mg CH_4_–C ha^−1^yr^−1^. This could be because the meta-analysis focused exclusively on the change from drained to rewetted peatlands, while in our case all the “conditions” are combinations of different habitat categories; also, the meta-analysis explored how the emissions change over time post-restoration, while we assumed immediate transition from one category (i.e. one set of EFs) to another, which can potentially include other ecological changes in addition to rewetting. Furthermore, the meta-analysis covers global studies, while Evans et al (2023), and hence our study, is based on Northern temperate peatlands as found in the UK.

There are some further caveats in considering how reliable our GHG estimates are likely to be. We assumed equal distributions of each habitat types in mosaic habitats, and some restoration works do not include a change between habitat types but only the proportion of each type. Thus, the actual differences between the restoration stages are probably larger than our estimation. Similarly, as we assumed the habitat types with the highest EFs when they were not clear from the descriptions provided, our estimations tend to be more conservative.

### Net GHG emissions and climate impacts over time

For most reporting purposes, total emissions (and removals) are expected to be communicated as GWP100 CO_2_-equivalent emissions. The large decrease in GWP100 net emissions after restoration, with a further reduction if further restored to ‘near-natural’ is a positive finding, and suggests incentives for organisations and countries to restore peatlands, for example in contributing to net-zero goals. In the context of the United Kingdom’s most recent Greenhouse Gas Inventory, our estimated emissions reduction from peatland restoration so far is equivalent to nearly 10% of the mitigation potential from Land-use, Land-use Change and Forestry (LULUCF) emissions till 2022 (Brown et al, 2024), which is a considerable amount. This highlights that peatland restoration is an important component for mitigation potential, as the largest emission sources from energy generation and transport will hopefully be addressed through wider decarbonisation, while the land sector will need its own set of emission reduction measures. Furthermore, there are concerns that these peatland restoration activities are not being sufficiently tracked and incorporated into national statistics (Brown, 2024), and our figure is only a portion of the total peatland restoration (and hence emissions reduction) for the UK, as it only covers data directly supplied through The Wildlife Trusts, and only from Trusts that had capacity to process and share their peatland restoration data for the purposes of this paper: additional peatland restoration is ongoing at Wildlife Trusts not captured here.

As highlighted in the introduction, however, GWP100 can fail to reveal the full climate impacts of emissions, and over the past decade, the usage of GWP100 has been suggested as a major limitation in understanding how peatland emissions affect climate (Neubauer, 2014; Neubauer & Megonigal, 2015). The UNEP (United Nations Environment Programme)-SETAC (Society of Environmental Toxicology and Chemistry) Life Cycle Initiative recommend reporting emissions using the 100-year Global Temperature change Potential (GTP100), to highlight longer-term impacts, and explore shorter-term impacts using the 20-year Global Warming Potential (GWP20) to help inform how effects can vary over time (Jolliet et al., 2018). Our results also showed a large reduction in GWP20 CO_2_e, indicating that there is still a substantial climate benefit from restoration in the near term, albeit less than in the longer-term perspective indicated by GWP100. An even greater reduction is reported under GTP100, with the suggested inference that the climate benefits of restoration are greater in the long-term: sufficiently great that the net CO_2_e footprint is negative, indicating the net effect of the CO_2_ removals and CH_4_ and N_2_O emissions is an overall cooling, rather than ‘less warming’, as seen in the shorter-term metrics. Furthermore, the long-term cooling effect can be nearly doubled if all sites are restored to the “target condition” (Table 3).

The GWP* results go further in helping illustrate what is happening over time, and how the different valuations merge from the range of pulse emission metrics. For the “initial condition” emissions associated with degraded peatlands, GWP* reveals that these cause continuously increasing warming for as long as the emissions are sustained. Over the first 20 years, both CH_4_ and CO_2_ build up and cause significant warming; but beyond this, the rate of increased warming from CH_4_ slows down, while CO_2_ increases at a fixed rate, dominating total warming over longer periods. After restoration, CH_4_-induced warming increases, especially for the initial couple of decades after the change in fluxes, but the CO_2_ removals mean that the growth of cumulative CO_2_-we emissions (i.e. increased warming from peatland emissions) is still lower than if left in a degraded state, and thus restoring is universally beneficial.

The two-step restoration offers an even greater climate benefit: whilst there is increased warming from CH_4_, again, especially in the first two decades after the CH_4_ emissions increase, the enhanced CO_2_ removals are enough that there is a decline in cumulative CO_2_we emissions: i.e., there is a year-on-year reduction in the increased temperatures caused by these peatlands. In the very long term, should these emission fluxes remain (CH_4_ and N_2_O continue to be emitted, but CO_2_ continues to be removed) total cumulative emissions would become neutral and then even negative. At this stage, even though the CH_4_ and N_2_O continue to cause warming, the CO_2_ removals are enough to [more than] compensate for the warming caused by emissions. This occurs even though annual GWP100 CO_2_e would still be positive in each individual year: because the sustained CH_4_ emissions do not actually cause ongoing cumulative increases in temperature each year, while the CO_2_ sequestration *does* lead to an ever-increasing contribution to cooling if removals continue. This cannot not be inferred from GWP100 reported emissions, hence there has been confusion over how northern peatlands with ‘positive net emissions’ can in fact have a net cooling effect (Frolking, Roulet & Fuglestvedt, 2006).

Our GWP* cumulative ‘CO_2_-warming-equivalent’ approach can highlight the same important principles as more complex methods. Günther et al. (2020) showed the climate impact of global peatland rewetting scenarios by modelling radiative forcing overtime. Although the scenario set-ups are different, they display similar dynamics and reach the same conclusions: in the absence of restoration, there is increasing, CO_2_-dominated warming from peatlands, whereas restoration has an immediate benefit in reducing warming, despite the warming effect of CH_4_ increases (see Figure 2 in Günther et al., 2020 and compare with our figure 1).

As it can indicate the temporal changes in temperature differing per gas, we suggest the GWP* cumulative CO_2_-we framework is a superior approach for exploring warming over time compared with the conventional pulse emission metrics. Other studies also highlight the weakness of the conventional GWP metrics in understanding ecosystem. Especially for wetland ecosystems, climate impacts have suggested the sustained-flux global warming potential (SGWP) as a superior alternative, given the expectation that habitats will continue to emit (Neubauer and Megonigal, 2015). GWP* can also reveal the impact of continuously sustained emissions, and has the advantage of further flexibility: the same simple equation can be used to report and understand the warming over time from any emission scenario, such as individual pulses, emissions sustained at the same rates, and any changes in the emission rate of interest. It is also a much simpler approach compared to fully modelling radiative forcing change over time (or resultant temperature impact) whilst still capturing the salient trends, as note above. For more robust atmospheric science purposes, however a more robust climate modelling approach should still be preferred.

Despite these advantages, GWP* does entail a fundamentally different approach to conventional reporting of CO_2_-equivalent footprints, requiring emission scenarios (even if based on straightforward assumptions of sustained emissions, as here) and interpretation that considers trends over time. If this is beyond the scope or purpose of a study, the 3 most common pulse metrics (GWP100, GTP100, GWP20) can still be used *together* to gain some further insight, as above.

No matter what methods are used for aggregating the total emissions, researchers working with systems involving multiple GHGs emissions are still recommended also to report on each gas individually, so that important details do not become irretrievably lost by aggregating as a CO_2_-equivalent. Then other emissions valuation or climate modelling approaches can still be explored by other users (Lynch, 2019).

Across all of our analyses, using both GWP* and the different pulse-metrics, a positive conclusion emerges: restoring peatlands results in less warming than if they remain degraded, with no risk of increased warming even in the short term, with a potential to have a long-term net cooling effect (although note discussion below further considering uncertainties in EFs).

### Biogeochemical drivers of the uncertainties in emission trajectories

The EFs calculated by Evans et al. (2023) have large uncertainty ranges, so our uncertainty analysis shows significant variations in total emissions (Figure 3), which corresponds with the global review by Darusman et al. (2023). Such uncertainties may derive from the temporal emissions change along the restoration process, as well as site-specific features or past management of the peatlands.

CO_2_ emissions/removals usually stabilise shortly after rewetting, while CH_4_ emissions undergo a significant increase over the first 4 years (Darusman et al., 2023). Surprisingly, a study measuring the GHG fluxes a decade after restoration showed a similar amount of CH_4_ being emitted as the unrestored site, except from the open water pools (Strack & Zuback, 2013). Since there is a lack of studies providing full continuous tracking of CH_4_ emissions over a long period post restoration, we cannot reliably picture the trajectory of CH_4_ emissions with restoration progress, even though sampling on methanogenic microbial community suggests CH_4_ flux is likely to stablise after 7-16 years (Urbanová & Bárta, 2020). In addition to the effects of changing (raising) water table depth, vegetation succession is also likely to influence CH_4_ fluxes as part of peatland restoration (Waddington & Day, 2007; Tuittila et al., 2000). Certain plant coverage, such as the species *Eriophorum vaginatum* (cotton grass), appear naturally as the water table rises and greatly increase CH_4_ emissions (Komulainen et al., 1998), as they create pathways for CH_4_ transportation, letting it release before being oxidised (Couwenberg & Fritz, 2012). On the contrary, *Sphagnum* mosses enhance CH_4_ cycling and reduce CH_4_ emissions (Larmola et al., 2010). The interacting effects of rewetting and changing plant communities make it challenging to narrow down EFs, particularly as these two important factors both change over time, potentially at different rates. Smyth et al. (2015), which is part of the data source for calculating the EFs by Evans et al. (2023), contains only the measurements from sites within 5 years after rewetting, during which the CH_4_ emissions are probably still under change, so the CH_4_ EFs have large uncertainty ranges, and may not be representative of the end-point emission.

It can take several years or decades for the peatlands to reach the emission levels suggested by the EFs (Kalhori et al., 2024; Wilson et al., 2016), and the time varies greatly among sites (Renou-Wilson et al., 2019). Since our estimation did not reflect the time required for habitat condition transitioning, and the total forcing only plateaus after rewetting (Günther et al., 2020), we suggest carrying out peatland restoration as early as possible for the climate mitigation purpose.

On the other hand, although restoration generally lowers CO_2_ and N_2_O emissions and increases CH_4_ emissions (Darusman et al., 2023), there are several individual cases showing unexpected effects contrary to these typical trends (Jauhiainen et al., 2008; Waddington, Strack & Greenwood, 2010). Waddington, Strack & Greenwood (2010) found that CO_2_ increased after rewetting due to an increase in plant production, which provides the organic matter sources for aerobic decomposition; while N_2_O emissions were highest, especially for bare peat, when the water table is rewetted to near surface, as Blondeau et al. (2024) found from lab incubation. Komulainen et al. (1998) measured CH_4_ emissions from rewetted sites to be lower than from pristine peatlands, but here are also examples where the CH_4_ emissions were very high after restoration and even more greatly overpowered the CO_2_ decrease, even when the site had been restored for more than 30 years (Vanselow-Algan et al., 2015). However, as the EFs used in our estimation were sourced from a sufficient variation of emissions measurements which reflect both inter- and intra-annual fluctuations, they are still likely to cover the CH_4_ spike shortly after rewetting. Our results strongly show that on a whole UK level restoration is likely to be beneficial. The worry of CH_4_ warming effect should not be the reason for delaying or cancelling restoration, as the long-lasting CO_2_ from degraded peatland causes continuous warming.

### Benefits beyond climate mitigation

Rewetting is only the first step of peatland restoration. Although it usually brings climate benefits, it rarely returns the species composition to natural states (Kreyling et al., 2021). Thus, further measures focusing on reintroducing the important species of pristine peatlands are necessary for restoring the rewetted peatlands to near-natural conditions. Our simulations with EF uncertainties show that there is a small possibility that total emissions will increase after changing the peatlands from “current” to “target” conditions, which mainly involves restoring from rewetted to near-natural states (Figure 3). Also, in our arbitrary O2R scenario, we indeed see some of the climate advantages being reversed by high CH4 emissions (Figure 1D). On the other hand, with the low risk of slightly reducing the climate benefit, restoring the peatlands to near-natural states can improve their biodiversity and resilience to extreme climate events (Strack et al., 2022; Deshmukh et al., 2021). Restoring peatlands gradually creates local island habitats that are more mild and humid than surrounding areas, as it changes microclimate conditions by changing the albedo and vegetation indices and reducing diurnal temperature fluctuation (Worrall et al., 2022). Intact peatlands have lower carbon loss in extreme drought than degraded peatlands (Deshmukh et al., 2021), and are likely to be more resilient to interannual weather changes than rewetted sites (Wilson et al., 2016). Moreover, with the increasing incidence of extreme weather events under climate change, drought and flooding will pose challenges in maintaining a constant water table, and could also lead to significant CO_2_, DOC emission and nitrogen runoff (Glatzel, Lemke & Gerold, 2006), so improving resilience should be an important focus in restoration.

The need to maximise climate mitigation and ecological services can in some cases be contradictory. Wilson et al. (2009) modelled that among 5 restoration scenarios, turning all peat-extraction sites to afforested peatlands provided the highest emissions savings, while restoring to near-natural conditions is most desirable for ecology. As nature-based solutions should also support, sustain, or enhance biodiversity (Seddon et al., 2020), emissions reduction should not be the only goal of peatland restoration.

### Further work and other considerations

Notably, our set-up for the O2R scenario is unlikely to happen in reality, as we selected the highest EFs for both CO_2_ and CH_4_, whereas their emissions are underpinned by water table depth in opposite directions. By treating the two gases independently for our study, we take a conservative approach that can reveal these more negative outlier cases. Further work could explore this by adapting the Monte-Carlo simulation approach we performed here by directly modelling interdependence of the EFs. For example, if an iteration selects a high EF for CH_4_, the range of possible values that can be chosen for CO_2_ is constrained, based on the equations linking water table depth, CH_4_ and CO_2_ emissions.

For other relevant future work, we reiterate our call above on the need for ongoing monitoring to confirm restoration is achievable and emissions do actually correspond with these EFs, and to control the water table as CH_4_ hotspots are possibly caused by inundation. Measuring the water table can also provide site-specific emission estimations using the equations from Evans et al. (2021), which is an easier way of generating more reliable estimations without flux towers. The more detailed emissions estimates should then also be explored beyond just the net GWP100 CO_2_e balance. Finally, there are also other restoration end points that are worth exploring.

Paludiculture is a possible alternative peatland management approach that can achieve emissions savings and provide both biodiversity and economical values, as it has been found in other European peatlands (Tanneberger et al., 2022), although there appear to be challenges in finding paludiculture practices that are both effective and practical in the UK context (Rhymes et al., 2023). The Wildlife Trusts are among the UK organisations currently trialing paludiculture projects.

## Conclusions

We have provided an in-depth exploration of the climate impacts of peatland from a realistic case-study, and using approaches which will be deployed for national reporting purposes. It is therefore a powerful finding that, within the context of the study, peatland restoration is almost universally positive. As would be expected from the relevant EFs, peatland restoration provides a significant climate benefit, especially in the initial rewetting of degraded peatlands. By comparing emissions across a range of emission metrics, and using GWP* to assess effects over time, we have explored most concerns about the initial strong warming impact where restoration increases CH_4_ emissions: even when accounting for this (and having demonstrated novel approaches to do so), restoration is still beneficial across all timeframes assuming typical EFs.

Considering the potential uncertainty ranges, further and more ecologically focussed restoration has a risk of reducing the climate benefit from the initially rewetted state, but mutually exclusive dependencies in peatland CH_4_ vs CO_2_ emissions make this unlikely in practice. Our simulation showed that even when the sites restored to “near-natural” conditions have high emissions of both gases, the impact in reducing the climate benefit would be minor, and even then still represents a substantial improvement over business as usual. Consequently, peatland restoration still presents a major contribution to climate change mitigation compared to leaving peatlands in a degraded state.

## Acknowledgements

We thank the staff at individual Wildlife Trusts for collating and sharing the data on peatland restoration used in this study, and the numerous site managers, wardens, contractors, project partners and volunteers for undertaking the peatland restoration activities that it documents.

---

**Supplementary Table S1.** The uncertainty statistics (mean, standard error and 95% confidence interval) of the total emissions of the 3 conditions under different metrics, calculated based on the Monte-Carlo simulation results.

| Total emissions<br>(t CO<sub>2</sub>e) | Mean | Standard<br>error | 5% CI | 95% CI |
| --- | --- | --- | --- | --- |
| Initial (GWP100) | 1063252.86 | 137.60 | 1062983.16 | 1063522.55 |
| Current (GWP100) | 79827.64 | 156.27 | 79521.35 | 80133.93 |
| Target (GWP100) | 32011.47 | 220.85 | 31578.59 | 32444.34 |
| Initial (GWP20) | 1267249.70 | 243.23 | 1266772.97 | 1267726.42 |
| Current (GWP20) | 409422.72 | 362.59 | 408712.05 | 410133.40 |
| Target (GWP20) | 373143.46 | 451.63 | 372258.27 | 374028.65 |
| Initial (GTP100) | 973657.37 | 117.36 | 973427.34 | 973887.41 |
| Current (GTP100) | -60331.12 | 104.99 | -60536.89 | -60125.35 |
| Target (GTP100) | -112377.04 | 123.31 | -112618.73 | -112135.35 |

## References

Abdalla, M., Hastings, A., Truu, J., Espenberg, M., Mander, Ü. & Smith, P. (2016) Emissions of methane from northern peatlands: a review of management impacts and implications for future management options. Ecology and Evolution. 6 (19), 7080–7102. doi:10.1002/ece3.2469.

Allen, M.R., Friedlingstein, P., Girardin, C.A.J., Jenkins, S., Malhi, Y., Mitchell-Larson, E., Peters, G.P. & Rajamani, L. (2022) Net Zero: Science, Origins, and Implications. Annual Review of Environment and Resources. 47 (1), 849–887. doi:10.1146/annurev-environ-112320-105050.

Allen, M.R., Fuglestvedt, J.S., Shine, K.P., Reisinger, A., Pierrehumbert, R.T. & Forster, P.M. (2016) New use of global warming potentials to compare cumulative and short-lived climate pollutants. Nature Climate Change. 6 (8), 773–776. doi:10.1038/nclimate2998.

Bao, T., Jia, G. & Xu, X. (2023) Weakening greenhouse gas sink of pristine wetlands under warming. Nature Climate Change. 13 (5), 462–469. doi:10.1038/s41558-023-01637-0.

Beyer, F., Jansen, F., Jurasinski, G., Koch, M., Schröder, B. & Koebsch, F. (2021) Drought years in peatland rewetting: rapid vegetation succession can maintain the net CO2 sink function. Biogeosciences. 18 (3), 917–935. doi:10.5194/bg-18-917-2021.

Blondeau, E., Velthof, G.L., Heinen, M., Hendriks, R.F.A., Stam, A., Van Den Akker, J.J.H., Weghorst, M. & Van Groenigen, J.W. (2024) Groundwater level effects on greenhouse gas emissions from undisturbed peat cores. Geoderma. 450, 117043. doi:10.1016/j.geoderma.2024.117043.

Bridgham, S.D., Megonigal, J.P., Keller, J.K., Bliss, N.B. & Trettin, C. (2006) The carbon balance of North American wetlands. Wetlands. 26 (4), 889–916. doi:10.1672/0277-5212(2006)26[889:TCBONA]2.0.CO;2.

Brown, K. (2024, August 9th) The Wildlife Trusts are restoring an enormous 60,000 hectares of peatlands – but it’s missing from UK Government’s net zero numbers. The Wildlife Trusts. https://www.wildlifetrusts.org/blog/kathryn-brown/wildlife-trusts-are-restoring-enormous-60000-hectares-peatlands-its-missing-uk

Brown, P., Cardenas, L., Del Vento, S., Karagianni, E., MacCarthy, J., Mullen, P., Gorji, S., Richmond, B., Thistlethwaite, G., Thomson, A., Wakeling, D. & Willis, D. (2024) UK Greenhouse Gas Inventory, 1990 to 2022: Annual Report for Submission under the Framework Convention on Climate Change.

Cain, M., Lynch, J., Allen, M.R., Fuglestvedt, J.S., Frame, D.J. & Macey, A.H. (2019) Improved calculation of warming-equivalent emissions for short-lived climate pollutants. npj Climate and Atmospheric Science. 2 (1), 29. doi:10.1038/s41612-019-0086-4.

Cherubini, F., Fuglestvedt, J., Gasser, T., Reisinger, A., Cavalett, O., Huijbregts, M.A.J., Johansson, D.J.A., Jørgensen, S.V., Raugei, M., Schivley, G., Strømman, A.H., Tanaka, K. & Levasseur, A. (2016) Bridging the gap between impact assessment methods and climate science. Environmental Science & Policy. 64, 129–140. doi:10.1016/j.envsci.2016.06.019.

Committee on Climate Change (2020) Land use: Policies for a Net Zero UK Committee on Climate Change. www.theccc.org.uk/publications.

Couwenberg, J. & Fritz, C. (2012) Towards developing IPCC methane ‘emission factors’ for peatlands (organic soils). Mires and Peat, Volume 10, Article 03, 1–17

Darusman, T., Murdiyarso, D., Impron & Anas, I. (2023) Effect of rewetting degraded peatlands on carbon fluxes: a meta-analysis. Mitigation and Adaptation Strategies for Global Change. 28 (3), 10. doi:10.1007/s11027-023-10046-9.

Deshmukh, C.S., Julius, D., Desai, A.R., Asyhari, A., Page, S.E., Nardi, N., Susanto, A.P., Nurholis, N., Hendrizal, M., Kurnianto, S., Suardiwerianto, Y., Salam, Y.W., Agus, F., Astiani, D., Sabiham, S., Gauci, V. & Evans, C.D. (2021) Conservation slows down emission increase from a tropical peatland in Indonesia. Nature Geoscience. 14 (7), 484–490. doi:10.1038/s41561-021-00785-2.

Dessus, B., Le Treut, H., & Laponche, B. (2008). Global warming: the significance of methane (INIS-FR--10-0531). France.

Drösler, M., Freibauer, A., Christensen, T.R. & Friborg, T. (2008) Observations and Status of Peatland Greenhouse Gas Emissions in Europe. In: A.J. Dolman, R. Valentini, & A. Freibauer (eds.). The Continental-Scale Greenhouse Gas Balance of Europe. Ecological Studies. New York, NY, Springer New York. pp. 243–261. doi:10.1007/978-0-387-76570-9_12.

Evans, C., Artz, R., Burden, A., Clilverd, H., Freeman, B., Heinemeyer, A., Lindsay, R., Morrison, R., Potts, J., Reed, M. & Williamson, J. (2023). Aligning the Peatland Code with the UK Peatland Inventory.

Evans, C., Artz, R., Moxley, J., Smyth, M-A., Taylor, E., Archer, N., Burden, A., Williamson, J., Donnelly, D., Thomson, A., Buys, G., Malcolm, H., Wilson, D., Renou-Wilson, F., Potts J. (2017). Implementation of an emission inventory for UK peatlands. Report to the Department for Business, Energy and Industrial Strategy, Centre for Ecology and Hydrology, Bangor. 88pp.

Evans, C.D., Peacock, M., Baird, A.J., Artz, R.R.E., Burden, A., et al. (2021) Overriding water table control on managed peatland greenhouse gas emissions. Nature. 593 (7860), 548–552. doi:10.1038/s41586-021-03523-1.

Flysjö, A., Henriksson, M., Cederberg, C., Ledgard, S. & Englund, J.-E. (2011) The impact of various parameters on the carbon footprint of milk production in New Zealand and Sweden. Agricultural Systems. 104 (6), 459–469. doi:10.1016/j.agsy.2011.03.003.

Frolking, S., Roulet, N. & Fuglestvedt, J. (2006) How northern peatlands influence the Earth’s radiative budget: Sustained methane emission versus sustained carbon sequestration. Journal of Geophysical Research: Biogeosciences. 111 (G1). doi:10.1029/2005JG000091.

Girardin, C.A.J., Jenkins, S., Seddon, N., Allen, M., Lewis, S.L., Wheeler, C.E., Griscom, B.W. & Malhi, Y. (2021) Nature-based solutions can help cool the planet-if we act now. Nature.593.

Glatzel, S., Lemke, S. & Gerold, G. (2006) Short-term effects of an exceptionally hot and dry summer on decomposition of surface peat in a restored temperate bog. European Journal of Soil Biology. 42 (4), 219–229. doi:10.1016/j.ejsobi.2006.03.003.

Günther, A., Barthelmes, A., Huth, V., Joosten, H., Jurasinski, G., Koebsch, F. & Couwenberg, J. (2020) Prompt rewetting of drained peatlands reduces climate warming despite methane emissions. Nature Communications. 11 (1), 1644. doi:10.1038/s41467-020-15499-z.

Harris, L.I., Richardson, K., Bona, K.A., Davidson, S.J., Finkelstein, S.A., Garneau, M., McLaughlin, J., Nwaishi, F., Olefeldt, D., Packalen, M., Roulet, N.T., Southee, F.M., Strack, M., Webster, K.L., Wilkinson, S.L. & Ray, J.C. (2022) The essential carbon service provided by northern peatlands. Frontiers in Ecology and the Environment. 20 (4), 222–230. doi:10.1002/fee.2437

Intergovernmental Panel On Climate Change (Ipcc) (2014) 2013 Supplement to the 2006 IPCC Guidelines for National Greenhouse Gas Inventories: Wetlands, Hiraishi, T., Krug, T., Tanabe, K., Srivastava, N., Baasansuren, J., Fukuda, M. and Troxler, T.G. (eds). Published: IPCC, Switzerland

Intergovernmental Panel On Climate Change (Ipcc) (2023) Climate Change 2021 – The Physical Science Basis: Working Group I Contribution to the Sixth Assessment Report of the Intergovernmental Panel on Climate Change. 1st edition. Cambridge University Press. doi:10.1017/9781009157896.

Jauhiainen, J., Limin, S., Silvennoinen, H. & Vasander, H. (2008) CARBON DIOXIDE AND METHANE FLUXES IN DRAINED TROPICAL PEAT BEFORE AND AFTER HYDROLOGICAL RESTORATION. Ecology. 89 (12), 3503–3514. doi:10.1890/07-2038.1.

Jolliet, O., Antón, A., Boulay, A.-M., Cherubini, F., Fantke, P., Levasseur, A., McKone, T.E., Michelsen, O., Milà I Canals, L., Motoshita, M., Pfister, S., Verones, F., Vigon, B. & Frischknecht, R. (2018) Global guidance on environmental life cycle impact assessment indicators: impacts of climate change, fine particulate matter formation, water consumption and land use. The International Journal of Life Cycle Assessment. 23 (11), 2189–2207. doi:10.1007/s11367-018-1443-y.

Kalhori, A., Wille, C., Gottschalk, P., Li, Z., Hashemi, J., Kemper, K. & Sachs, T. (2024) Temporally dynamic carbon dioxide and methane emission factors for rewetted peatlands. Communications Earth & Environment. 5 (1), 62. doi:10.1038/s43247-024-01226-9.

Komulainen, V.-M., Nykänen, H., Martikainen, P.J. & Laine, J. (1998) Short-term effect of restoration on vegetation change and methane emissions from peatlands drained for forestry in southern Finland. Canadian Journal of Forest Research. 28 (3), 402–411. doi:10.1139/x98-011.

Kreyling, J., Tanneberger, F., Jansen, F., Van Der Linden, S., Aggenbach, C., et al. (2021) Rewetting does not return drained fen peatlands to their old selves. Nature Communications. 12 (1), 5693. doi:10.1038/s41467-021-25619-y.

Larmola, T., Tuittila, E.-S., Tiirola, M., Nykänen, H., Martikainen, P.J., Yrjälä, K., Tuomivirta, T. & Fritze, H. (2010) The role of Sphagnum mosses in the methane cycling of a boreal mire. Ecology. 91 (8), 2356–2365. doi:10.1890/09-1343.1.

Li, T., Canadell, J.G., Yang, X.-Q., Zhai, P., Chao, Q., Lu, Y., Huang, D., Sun, W. & Qin, Z. (2022) Methane Emissions from Wetlands in China and Their Climate Feedbacks in the 21st Century. Environmental Science & Technology. 56 (17), 12024–12035. doi:10.1021/acs.est.2c01575.

Lynch, J. (2019) Availability of disaggregated greenhouse gas emissions from beef cattle production: A systematic review. Environmental Impact Assessment Review. 76, 69–78. doi:10.1016/j.eiar.2019.02.003.

Lynch, J., Cain, M., Pierrehumbert, R. & Allen, M. (2020) Demonstrating GWP*: a means of reporting warming-equivalent emissions that captures the contrasting impacts of short- and long-lived climate pollutants. Environmental Research Letters. 15 (4), 044023. doi:10.1088/1748-9326/ab6d7e.

Ma, S., Creed, I.F. & Badiou, P. (2024) New perspectives on temperate inland wetlands as natural climate solutions under different CO2-equivalent metrics. npj Climate and Atmospheric Science. 7 (1), 222. doi:10.1038/s41612-024-00778-z.

Martikainen, P.J., Nykänen, H., Crill, P. & Silvola, J. (1993) Effect of a lowered water table on nitrous oxide fluxes from northern peatlands. Nature. 366 (6450), 51–53. doi:10.1038/366051a0.

McAuliffe, G.A., Lynch, J., Cain, M., Buckingham, S., Rees, R.M., Collins, A.L., Allen, M., Pierrehumbert, R., Lee, M.R.F. & Takahashi, T. (2023) Are single global warming potential impact assessments adequate for carbon footprints of agri-food systems? Environmental Research Letters. 18 (8), 084014. doi:10.1088/1748-9326/ace204.

Milne, A.E., Glendining, M.J., Bellamy, P., Misselbrook, T., Gilhespy, S., Rivas Casado, M., Hulin, A., Van Oijen, M. & Whitmore, A.P. (2014) Analysis of uncertainties in the estimates of nitrous oxide and methane emissions in the UK’s greenhouse gas inventory for agriculture. Atmospheric Environment. 82, 94–105. doi:10.1016/j.atmosenv.2013.10.012.

Neubauer, S.C. (2014) On the challenges of modeling the net radiative forcing of wetlands: reconsidering Mitsch et al. 2013. Landscape Ecology. 29 (4), 571–577. doi:10.1007/s10980-014-9986-1.

Neubauer, S.C. & Megonigal, J.P. (2015) Moving Beyond Global Warming Potentials to Quantify the Climatic Role of Ecosystems. Ecosystems. 18 (6), 1000–1013. doi:10.1007/s10021-015-9879-4.

Page, S.E. & Baird, A.J. (2016) Peatlands and Global Change: Response and Resilience. Annual Review of Environment and Resources. 41 (1), 35–57. doi:10.1146/annurev-environ-110615-085520.

Peacock, M., Audet, J., Bastviken, D., Futter, M.N., Gauci, V., Grinham, A., Harrison, J.A., Kent, M.S., Kosten, S., Lovelock, C.E., Veraart, A.J. & Evans, C.D. (2021) Global importance of methane emissions from drainage ditches and canals. Environmental Research Letters. 16 (4), 044010. doi:10.1088/1748-9326/abeb36.

Pierrehumbert, R.T. (2014) Short-Lived Climate Pollution. Annual Review of Earth and Planetary Sciences. 42 (1), 341–379. doi:10.1146/annurev-earth-060313-054843.

Qiu, C., Ciais, P., Zhu, D., Guenet, B., Chang, J., et al. (2022) A strong mitigation scenario maintains climate neutrality of northern peatlands. One Earth. 5 (1), 86–97. doi:10.1016/j.oneear.2021.12.008.

R Core Team (2023). _R: A Language and Environment for Statistical Computing_. R. Foundation for Statistical Computing, Vienna, Austria. <https://www.R-project.org/>.

Renou-Wilson, F., Moser, G., Fallon, D., Farrell, C.A., Müller, C. & Wilson, D. (2019) Rewetting degraded peatlands for climate and biodiversity benefits: Results from two raised bogs. Ecological Engineering. 127, 547–560. doi:10.1016/j.ecoleng.2018.02.014.

Rhymes, J.M., Arnott, D., Chadwick, D.R., Evans, C.D. & Jones, D.L. (2023) Assessing the effectiveness, practicality and cost effectiveness of mitigation measures to reduce greenhouse gas emissions from intensively cultivated peatlands. Land Use Policy. 134, 106886. doi:10.1016/j.landusepol.2023.106886.

Seddon, N., Chausson, A., Berry, P., Girardin, C.A.J., Smith, A. & Turner, B. (2020) Understanding the value and limits of nature-based solutions to climate change and other global challenges. Philosophical Transactions of the Royal Society B: Biological Sciences. 375 (1794), 20190120. doi:10.1098/rstb.2019.0120.

Smith, M.A., Cain, M. & Allen, M.R. (2021) Further improvement of warming-equivalent emissions calculation. npj Climate and Atmospheric Science. 4 (1), 19. doi:10.1038/s41612-021-00169-8.

Smyth, M.A., Taylor, E.S., Birnie, R.V., Artz, R.R.E., Dickie, I., Evans, C., Gray, A., Moxey, A., Prior, S., Littlewood, N. and Bonaventura, M. (2015) Developing Peatland Carbon Metrics and Financial Modelling to Inform the Pilot Phase UK Peatland Code. Report to Defra for Project NR0165, Crichton Carbon Centre, Dumfries.

Strack, M., Davidson, S.J., Hirano, T. & Dunn, C. (2022) The Potential of Peatlands as Nature-Based Climate Solutions. Current Climate Change Reports. 8 (3), 71–82. doi:10.1007/s40641-022-00183-9.

Strack, M. & Zuback, Y.C.A. (2013) Annual carbon balance of a peatland 10 yr following restoration. Biogeosciences. 10 (5), 2885–2896. doi:10.5194/bg-10-2885-2013.

Tanneberger, F., Birr, F., Couwenberg, J., Kaiser, M., Luthardt, V., Nerger, M., Pfister, S., Oppermann, R., Zeitz, J., Beyer, C., Van Der Linden, S., Wichtmann, W. & Närmann, F. (2022) Saving soil carbon, greenhouse gas emissions, biodiversity and the economy: paludiculture as sustainable land use option in German fen peatlands. Regional Environmental Change. 22 (2), 69. doi:10.1007/s10113-022-01900-8.

Trenbirth, H. & Dutton, A. (2019). UK natural capital: peatlands. London, UK: Office for National Statistics. See https://www.ons.gov.uk.

Tuittila, E., Komulainen, V., Vasander, H., Nykänen, H., Martikainen, P.J. & Laine, J. (2000) Methane dynamics of a restored cut-away peatland. Global Change Biology. 6 (5), 569–581. doi:10.1046/j.1365-2486.2000.00341.x.

Urbanová, Z. & Bárta, J. (2020) Recovery of methanogenic community and its activity in long-term drained peatlands after rewetting. Ecological Engineering. 150, 105852. doi:10.1016/j.ecoleng.2020.105852.

Vanselow-Algan, M., Schmidt, S.R., Greven, M., Fiencke, C., Kutzbach, L. & Pfeiffer, E.-M. (2015) High methane emissions dominated annual greenhouse gas balances 30 years after bog rewetting. Biogeosciences. 12 (14), 4361–4371. doi:10.5194/bg-12-4361-2015.

Waddington, J.M. & Day, S.M. (2007) Methane emissions from a peatland following restoration. Journal of Geophysical Research: Biogeosciences. 112 (G3), 2007JG000400. doi:10.1029/2007JG000400.

Waddington, J.M., Strack, M. & Greenwood, M.J. (2010) Toward restoring the net carbon sink function of degraded peatlands: Short-term response in CO2 exchange to ecosystem-scale restoration. Journal of Geophysical Research: Biogeosciences. 115 (G1). doi:10.1029/2009JG001090.

Wilson, D., Alm, J., Laine, J., Byrne, K.A., Farrell, E.P. & Tuittila, E. (2009) Rewetting of Cutaway Peatlands: Are We Re-Creating Hot Spots of Methane Emissions? Restoration Ecology. 17 (6), 796–806. doi:10.1111/j.1526-100X.2008.00416.x.

Wilson, D., Farrell, C.A., Fallon, D., Moser, G., Müller, C. & Renou-Wilson, F. (2016) Multiyear greenhouse gas balances at a rewetted temperate peatland. Global Change Biology. 22 (12), 4080–4095. doi:10.1111/gcb.13325.

Worrall, F., Howden, N.J.K., Burt, T.P., Rico-Ramirez, M.A. & Kohler, T. (2022) Local climate impacts from ongoing restoration of a peatland. Hydrological Processes. 36 (3), e14496. doi:10.1002/hyp.14496.

Young, H., Nickerson, R., Clilverd, H., Buys, G., Tomlinson, S., Thomson, A. & Henshall, P. (2023). Mapping greenhouse gas emissions & removals for the land use, land-use change & forestry sector. A report of the National Atmospheric Emissions Inventory 1990-2021.London, Department for Energy Security and Net Zero, 56pp

